# EnhancerP-2L: A Gene regulatory site identification tool for DNA enhancer region using CREs motifs

**DOI:** 10.1101/2020.01.20.912451

**Authors:** Ahmad Hassan Butt, Salem Alkhalaf, Shaukat Iqbal, Yaser Daanial Khan

## Abstract

Enhancers are DNA fragments that do not encode RNA molecules and proteins, but they act critically in the production of RNAs and proteins by controlling gene expression. Prediction of enhancers and their strength plays significant role in regulating gene expression. Prediction of enhancer regions, in sequences of DNA, is considered a difficult task due to the fact that they are not close to the target gene, have less common motifs and are mostly tissue/cell specific. In recent past, several bioinformatics tools were developed to discriminate enhancers from other regulatory elements and to identify their strengths as well. However the need for improvement in the quality of its prediction method requires enhancements in its application value practically. In this study, we proposed a new method that builds on nucleotide composition and statistical moment based features to distinguish between enhancers and non-enhancers and additionally determine their strength. Our proposed method achieved accuracy better than current state-of-the-art methods using 5-fold and 10-fold cross-validation. The outcomes from our proposed method suggest that the use of statistical moments based features could bear more efficient and effective results. For the accessibility of the scientific community, we have developed a user-friendly web server for EnhancerP-2L which will increase the impact of bioinformatics on medicinal chemistry and drive medical science into an unprecedented resolution. Web server is freely accessible at http://www.biopred.org/enpred.

## 1. Introduction

In cellular biology, regulation of transcription is performed to recruit elongation factors or RNA polymerase II initiation. This is mainly achieved at specific sequences of DNA by binding transcriptional factors (TFs). Transcription initiation sites are harbored by promoter regions which are the most studied sites in DNA [1]. Some DNA sequences have multiple transcription factor binding sites and are near or far away from promoter regions. Such DNA segments are denoted as enhancers [2], [3]. The transcription of genes is boosted by enhancers which influence various cellular activities such as cell carcinogenesis and virus activity, tissue specificity of gene expression, differentiation and cell growth, regulation and gene expression and develop relationship between such processes very closely [4].

Enhancers can be a short (50-1500bp) segment of DNA and situated 1Mbp (1,000,000bp) distance away from a gene. Sometimes they can even exist in different chromosomes [5], [6]. On the other hand, promoters are located near the start of the transcription sites of a gene. Due to this fact of locational difference between promoters and enhancers, the task of enhancer’s prediction is highly difficult and challenging than promoters [7]. Many human diseases like inflammatory bowel disease, disorder and various cancers have been linked to this genetic variation in enhancers [8]–[11].

About 30 years ago, a DNA segment, which was the first characterized enhancer, increased the transcription of β-globin gene during a transgenic assay inside the virus genome of SV40 tumor [12]. Scientific research during recent past has discovered that enhancers have many subgroups such as weak and strong enhancers, latent enhancers and poised enhancers [13]. Prediction of enhancers and their subgroups is an interesting area of research as they are considered important in disease and evolution. In higher classification of eukaryotes, transcription factor repertoire, diverse in nature, binds to enhancers [14]. This process of binding orchestrates many cellular events that are critical to the cellular system. Some of the events that are coordinated through this binding are maintenance of the cell identity, differentiation and response to stimuli [15], [16].

In the past, purely experimental techniques were being relied upon for the prediction of enhancers. Pioneering works in enhancer prediction was proposed in [4], [17]. The former was to use combinations such as transcription factor, P300 [18], with enhancers to identify them. This method would usually under-detect or miss the concerned targets and resulted in high failure rates due to the fact that all enhancers do not have transcription factor occupations. The latter was to utilize DNase I hypersensitivity for enhancer predictions. Hence, leading to high false positive rate as many other DNA segments, which were non-enhancers, detected incorrectly as enhancers. Although, genome-wide mapping techniques of histone modifications [1], [19]–[23] could improve the aforesaid deficiencies in the prediction of promoters and enhancers, but they are time consuming and expensive.

Several bioinformatics tools have been developed for rapid and cost effective classification of enhancers in genomics. CSI-ANN [21] used data transformations efficiently to formulate the samples and predict using Artificial-Neural-Network (ANN) classifications. EnhancerFinder [1] incorporated evolutionary conservation information features into sample formulation combined with a multiple kernel learning algorithm as a classifier. RFECS [23] applied random forest algorithm for improvements in detection methods. EnhancerDBN [24] used deep belief networks for enhancer predictions. BiRen [25] increased the predictive performance by utilizing deep learning based method. By utilizing these bioinformatics tools, enhancer detection can be achieved by the research community. Formed using many different large sub-groups of functional elements, enhancers can be grouped as weak, strong, inactive and poised enhancers. iEnhancer-2L [26], the first ever predictor to detect enhancers and identify their strengths and was based on sequence information only. Pseudo K-tuple nucleotide compositions (PseKNC) based features were incorporated into iEnhancer-2L. It has been used in many analysis related to genomics increasingly. Furthermore, many other methods, such as EnhancerPred [27] and EnhancerPred_2.0 [28], were introduced to improve the performance by incorporating other features based on DNA sequences. iEnhancer-5Step [29] was recently developed using the hidden information of DNA sequences infused with Support Vector Machine (SVM) based predictions. However, improvement in the performance of the aforementioned predictors is still required. Specifically, the success rate of discriminating strong and weak enhancers is not up to the expectations of the scientific community. The current study is initiated to propose a method which would deal with this problem.

## 2. Materials and Methods

### 2.1 Benchmark Dataset

The benchmark dataset of DNA enhancer sites, originally constructed and used in recent past [26], was re-used in the proposed method. In the current dataset, information related to nine different cell lines (K562, H1ES, HepG2, GM12878, HSMM, HUVEC, NHEK, NHLF and HMEC) was used in the collection of enhancers and 200bp fragments were extracted from DNA sequences. To remove pairwise sequences from the dataset, CD-HIT [30] tool was used to remove sequences having more than 20% similarity. The benchmark dataset includes 2,968 DNA enhancer sequences from which 1,484 are non-enhancer sequences and 1,484 are enhancer sequences. From 1,484 enhancer sequences, 742 are strong enhancers and 742 are weak enhancers for the second layer classification. Furthermore, the independent dataset used by a recent study [29] was utilized to enhance the effectiveness and performance of the proposed model. The independent dataset included 400 DNA enhancer sequences from which 200 (100 strong and 100 weak enhancers) are enhancers and 200 are non-enhancers. Table.1 and Table.2 includes the breakdown of the benchmark dataset.

In statistical based prediction models, the benchmark dataset mostly include training datasets and testing datasets. By utilizing various benchmark datasets, results obtained are computed from 5-fold and 10-fold cross-validations. The definition of a benchmark dataset is used in eqs.(1):

**Table: 1.** Breakdown of the benchmark datasets of DNA Enhancers and Non-Enhancers.

| DNA Samples | Benchmark Dataset [26] | Independent Dataset [29] |
| --- | --- | --- |
| Non-Enhancers | 1,484 | 200 |
| Enhancers | 1,484 | 200 |
| Overall | 2,968 | 400 |

**Table: 2.** Breakdown of the DNA Enhancers from benchmark datasets.

| DNA Enhancers | Benchmark Dataset [26] | Independent Dataset [29] |
| --- | --- | --- |
| Strong Enhancers | 742 | 100 |
| Weak Enhancers | 742 | 100 |
| Total Enhancer | 1,484 | 200 |

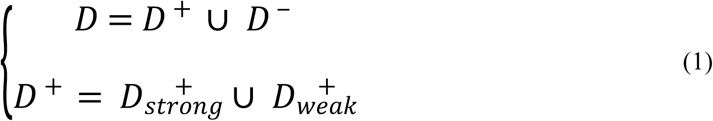

Where *D* ^+^ contains 1,484 enhancers and *D* ^−^ contains 1,484 non-enhancers. 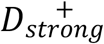 contains 742 strong enhancers, 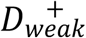 contains 742 weak enhancers and U denotes the symbol of “union” in the set theory.

### 2.2 Feature Extraction

Today, one of the core requirements in the development of an effective bioinformatics predictor is the need to formulate the biological sequence with a vector or a discrete model without losing any key-order characteristics or sequence-pattern information. The reason for this fact, as explained in a comprehensive state-of-the-art review [31], that the existing machine-learning algorithms cannot handle sequences directly but rather in vector formulations. However, there exists some possibility that all the sequence-pattern information from a vector might be lost in a discrete model formulation. To overcome the sequence-pattern information loss from proteins, Chou proposed pseudo amino acid composition (PseAAC) [32]. In almost all areas of bioinformatics and computational proteomics [31], the Chou’s PseAAC concept has been widely used ever since it was proposed. In the recent past, three publicly accessible and powerful softwares, ‘propy’ [33], ‘PseAAC-Builder’ [34] and ‘PseAAC-General’ [35] were developed and the importance and popularity of Chou’s PseAAC in computational proteomics has increased more ever since. ‘PseAAC-General’ calculates Chou’s general PseAAC [36] and the other two softwares generate Chou’s special PseAAC in various modes [37]. The Chou’s general PseAAC included not only the feature vectors of all the special modes, but also the feature vectors of higher levels, such as “Gene Ontology” mode [36], “Functional Domain” mode [36] and “Sequential Evolution” mode or “PSSM” mode [36]. Using PseAAC successfully for finding solutions to various problems relevant to peptide/protein sequences, encouraged the idea to introduce PseKNC (Pseudo K-tuple Nucleotide Composition) [38] for generating different feature vectors for DNA/RNA sequences [39], [40] which proved very effective and efficient as well. In recent times a useful, efficient and a very powerful web server called ‘Pse-in-One’ [41] and its recently updated version ‘Pse-in-One2.0’ [42] were developed that are able to generate any preferred feature vector of pseudo components for DNA/RNA and protein/peptide sequences according to the requirements of study by any researcher.

In this study, we utilized the Kmer [43] approach to represent the DNA sequences. According to Kmer, the occurrence frequency of ‘n’ neighboring nucleic acids can be represented from a DNA sequence. Hence, by using the sequential model, a sample of DNA having ‘w’ nucleotides is expressed generally as eqs.(2)

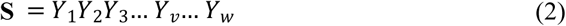

Where *Y*_1_ is represented as the first nucleotide of the DNA sample **S**, *Y*_2_ as the second nucleotide having the 2^nd^ position of occurrence in DNA sample S and so on so fourth *Y*_*w*_ denotes the last nucleotide of the DNA sample. ‘w’ is the total length of the nucleotides in a DNA sample. The *Y*_*v*_nucleotide can be any four of the nucleotides which can be represented using the aforementioned discrete model. The nucleotide *Y*_*v*_can be further described using eqs. (3)

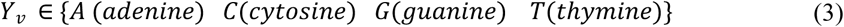

Here ∈ is the symbol used to represent the set theory ‘member of’ property and 1 ≤ *v* ≤ *n*. The components that are defined by the aforementioned discrete model utilize relevant nucleotides useful features to expedite the extraction methods. These components are further used in statistical moments based feature extraction methods.

#### 2.2.1 Statistical Moments

Statistical moments are certain types of quantitative measures that are used for the study of the concentrations of some key configurations in a collection of data used for pattern recognition related problems [44]. Several properties of data are described by different orders of moments. Some moments are used to reveal eccentricity and orientation of data while some are used to estimate the data size [45]–[48]. Several moments have been formed by various mathematicians and statisticians based on famous distribution functions and polynomials. To explicate the current problem, Raw; Central and Hahn moments are utilized [49].

The moments that are used in calculations of mean, variance and asymmetry of the probability distribution are known as raw moments. They are neither location-invariant nor scale-invariant. Similar type of information is obtained from the Central moments, but these central moments are calculated using the centroid of the data. The central moments are location-invariant with respect to centroid as they are calculated along the centroid of the data, but still they remain scale-variant. The moments based on Hahn polynomials are known as Hahn moments. These moments are neither location-variant nor scale-invariant [50]–[53]. The fact that these moments are sensitive to biological sequence ordered information amplifies the reason to choose them as they are primarily significant in extracting the obscure features from DNA sequences. The use of scale-invariant moment has consequently been avoided during the current study. The values quantified from utilizing each method enumerate data on its own measures. Furthermore, the variations in data source characteristics imply variations in the quantified value of moments calculated for arbitrary datasets. In the current study, the 2D version of the aforementioned moments is used and hence the linear structured DNA sequence as expressed by eqs. (2) is transformed into a 2D notation. The DNA sequence, which is 1D, is transformed to a 2D structure using row major scheme through the following eqs. (4):

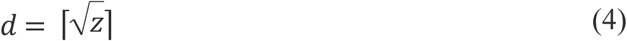

Where the sample sequence length is ‘z’ and the 2-dimensional square matrix has ‘*d*’ as its dimension. The ordering obtained from equation (4) is used to form matrix M (eqs.(5)) having ‘m’ rows and ‘m’ columns.

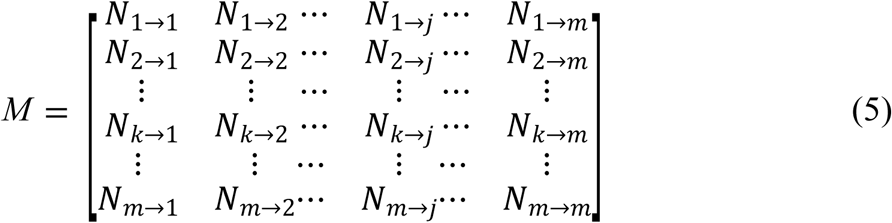

The transformation from M matrix to square matrix M’ is performed using the mapping function ‘?’. This function is defined as eqs.(6):

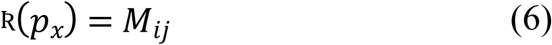

If the population of square matrix M’ is done as row major order then, 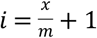 and *j* = *x mod m*.

Any vector or matrix, which represents any pattern, can be used to compute different forms of moments. The values of M’ are used to compute raw moments. The moments of a 2D continuous function *A*(*j, k*) to order (j + k) are calculated from eqs.(7):

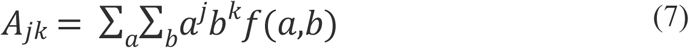

The raw moments of 2D matrix M, with order (j + k) and up to a degree of 3, are computed using the eqs(7). The origin of data as the reference point and distant components from the origin are assumed and utilized by the raw moments for computations. The 10 moment features computed up to degree-3 are labeled as *M*_00_,*M*_01_, *M*_10_, *M*_11_, *M*_02_, *M*_20_, *M*_12_, *M*_21_, *M*_30_ & *M*_03_.

The centroid of any data is considered as its center of gravity. The centroid is the point in the data where it is uniformly distributed in all directions in the relations of its weighted average [54], [55]. The central moments are also computed up to degree-3, using the centroid of the data as their reference point, from the following eqs.(8):

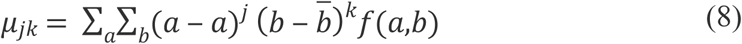

The degree-3 central moments with 10 distinct features are labeled as *μ*_00_, *μ*_01_, *μ*_10_, *μ*_11_, *μ*_02_, *μ*_20_, *μ*_12_, *μ*_21_, *μ*_30_ & *μ*_03_. The centroids *a* and 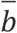 are calculated from eqs.(9) and eqs.(10):

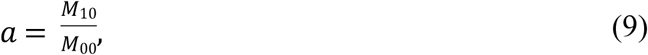

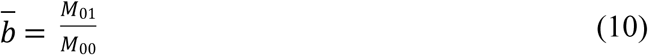

The Hahn moments are computed by transforming 1D notations into square matrix notations. This square matrix is valuable for the computations of discrete Hahn moments or orthogonal moments as these moments are of 2D form and require a two-dimensional square matrix as input data. These Hahn moments are orthogonal in nature that implies that they possess reversible properties. Usage of this property enables the reconstruction of the original data using the inverse functions of discrete Hahn moments. This further indicates that the compositional and positional features of a DNA sequence are somehow conserved within the calculated moments. M’ matrix is used as 2D input data for the computations of Orthogonal Hahn moments. The order ‘m’ Hahn polynomial can be computed from eqs.(11):

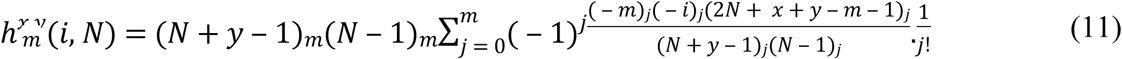

The aforementioned Pochhammer symbol 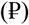 was defined as follows in eqs.(12):

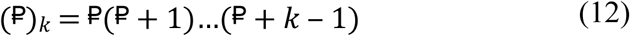

And was simplified further by the Gamma operator in eqs.(13):

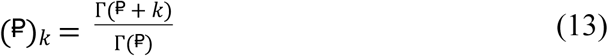

The Hahn moments raw values are scaled using a weighting function and a square norm given as in eqs.(14):

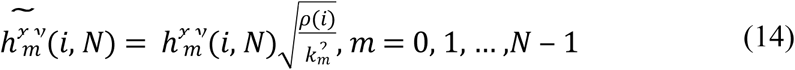

Meanwhile, in eqs.(15),

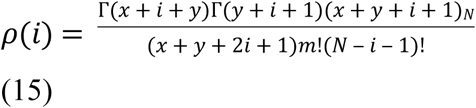

The Hahn moments are computed up to degree-3 for the 2-D discrete data as follows in eqs.(16):

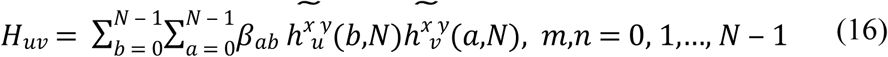

The 10 key Hahn moments based features are represented by *H*_00_, *H*_01_, *H*_10_, *H*_11_, *H*_02_, *H*_20_, *H*_12_, *H*_21_, *H*_30_ & *H*_03_. Matrix M’ was utilized in computing 10 Raw, 10 Central and 10 Hahn moments for every DNA sample sequence up to degree-3 which later are unified into the miscellany Super Feature Vector (SFV).

#### 2.2.2 DNA-Position-Relative-Incident-Matrix (D-PRIM)

The DNA characteristics such as ordered location of the nucleotides in the DNA sequences are of pivotal significance for identification. The relative positioning of nucleotides in any DNA sequence is considered core patterns prevailing the physical features of the DNA sequence. The DNA sequence is represented by D-PRIM in (4 × 4) order. The matrix in eqs.(17) is used to extract position-relative attributes of every nucleotide in the given DNA sequence.

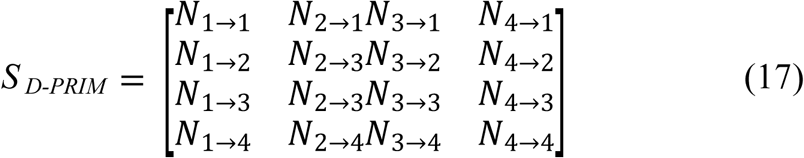

Here the indication score of the *y*^th^ position nucleotide is determined using *N*_*x*→*y*_ with respect to the x^th^ nucleotide first occurrence in the sequence. The nucleotide type ‘*y*’ substitutes this score in the biological evolutionary process. The occurrence positional values, in alphabetical order, represented as 4 native nucleotides. The S _D-PRIM_ matrix is formed with 16 coefficient values obtained after successfully performing computations on position relative incidences.

30 raw, central and Hahn moments (10 raw, 10 central & 10 Hahn), up to degree-3, were computed using the 2D S _D-PRIM_ matrix through which 30 features were obtained with 16 unique coefficients and were further incorporated into the miscellany Super Feature Vector (SFV).

#### 2.2.3 DNA-Reverse-Position-Relative-Incident-Matrix (D-RPRIM)

It often happens in cellular biology that the same ancestor is responsible for evolving more than one DNA sequence. These cases mostly outcome homologous sequences. The performance of the classifier is hugely effected by these homologous sequences and hence for producing accurate results, sequence similarity searching is reliable and effectively useful. In machine learning, accuracy and efficiency is hugely dependent on the meticulousness and thoroughness of algorithms through which most pertinent features in the data are extracted. The algorithms used in machine learning have the ability to learn and adapt the most obscure patterns embedded in the data while understanding and uncovering them during the learning phase. The procedure followed during the computation of D-PRIM was utilized in computations of D-RPRIM but only with reverse DNA sequence ordering. This procedure further uncovered hidden patterns for prediction and ambiguities between similar DNA sequences were also alleviated. The 2D matrix D-RPRIM was formed with (4 × 4) order having 16 unique coefficients. It is defined by eqs.(18):

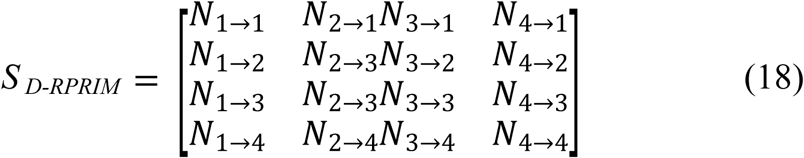

Similarly, 30 raw, central and Hahn moments (10 raw, 10 central & 10 Hahn), up to degree-3, were computed using the 2D S _D-RPRIM_ matrix through which 30 features were also obtained with 16 unique coefficients and they were also incorporated into the miscellany Super Feature Vector (SFV).

#### 2.2.4 Frequency-Distribution-Vector (FDV)

The distribution of occurrence of every nucleotide was used to compute the frequency distribution vector. The frequency distribution vector (FDV) is defined as in eqs.(19):

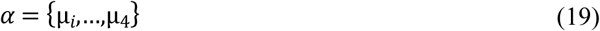

Here µ_*i*_is the frequency of occurrence of the *i*^th^ (1 ≤ *i* ≤ 4) relevant nucleotide. Furthermore, the relative positions of nucleotides in any sequence is alleviated using these measures. The miscellany Super Feature Vector (SFV) includes these 4 features from FDV as unique attributes.

#### 2.2.6 D-AAPIV (DNA-Accumulative-Absolute-Position-Incidence-Vector)

The distributional information of nucleotides was stored using frequency distribution vector which used the obscure features of DNA sequences in relevance to the compositional details. FDV does not have any information regarding relative positional details of relevant nucleotide residues in DNA sequences. This relative positional information was accommodated using D-AAPIV with a length of 4 critical features associated with 4 native nucleotides in a DNA sequence. These 4 critical features from D-AAPIV are also added into the miscellany Super Feature Vector (SFV).

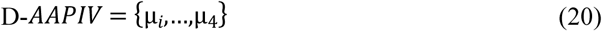

Here µ_*i*_is any element of D-AAPIV, from DNA sequence *S*_*j*_ having ‘n’ total nucleotides, which can be calculated using eqs.(21):

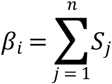

#### 2.2.7 D-RAAPIV (DNA-Reverse-Accumulative-Absolute-Position-Incidence-Vector)

D-RAAPIV is calculated using the reverse DNA sequence as input with the same method used using D-AAPIV calculations. This vector is calculated to find the deep and hidden features of every sample with respect to reverse relative positional information. D-RAAPIV is formed as the following eqs.(24) using the reversed DNA sequence and generates 4 valuable features. These 4 critical features from D-RAAPIV are also added into the miscellany Super Feature Vector (SFV).

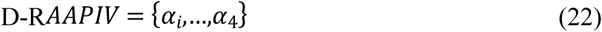

Here *α*_*i*_is any element of D-RAAPIV, from DNA sequence *S*_*j*_ having ‘n’ total nucleotides, which can be calculated using eqs.(23):

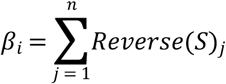

After calculating all possible features from the aforementioned extraction methods, the Super Feature Vector (SFV) of 134D features was obtained, for further processing in classification algorithm.

### 2.3 Classification Algorithm

#### 2.3.1 Random Forests

In the past, ensemble learning methods have been applied in many bioinformatics relevant research studies [56], [57] and have produced highly efficient outcomes in measures of performance. Ensemble learning methods utilize many classifiers in a classification problem with aggregation of their results. The two most commonly used methods are boosting [58], [59] and bagging [60] which perform classifications using trees. In boosting, the trees which are successive, propagate extra weights to points which are predicted incorrectly by the previous classifiers. The weighted vote decides the prediction in the end. Whereas, in bagging, the successive trees do not rely on previous trees, rather, each tree is constructed independently from the data using a bootstrap sample. The simple majority vote decides the prediction in the end.

Leo Breiman [61] built random forests, through which an additional layer of randomness is added to bagging. The random forests changed the construction of the classification trees by adding the construction of each tree from the data using a different bootstrap sample. The splitting of each node, in standard classification trees, is performed by dividing each node equally among all the variables. However, in random forests, the splitting of each node is performed by choosing the best among a subset of predictors which are chosen randomly at that node (Figure 1 shows the structure of the random forest classifier). As compared to many other classifiers, such as support vector machine, discriminant analysis and neural networks, this counterintuitive strategy perform very well and is robust against overfitting [56].

**Fig.1:**
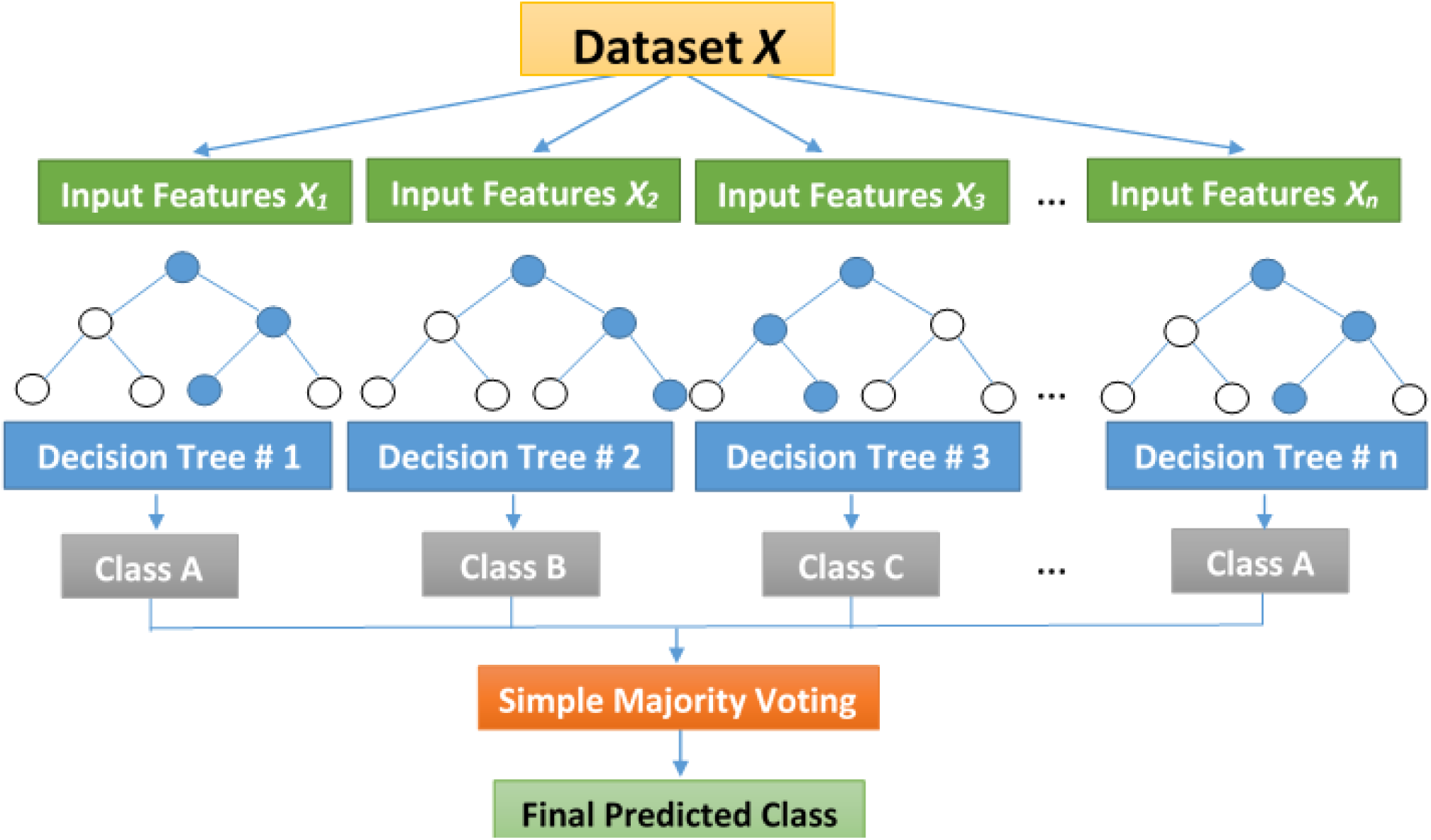
The structure of the Random Forest classifier.

##### 2.3.1.1 Algorithm: Supervised learning using Random Forest

Scikit-Learn [62] library using python was implemented for random forest classifier for fitting the trainings and simulations in our proposed method. The number of trees was increased from the default parameter value of 10 to 25. According to [63], 128 is the theoretical upper limit of the trees and any further increase in the number of trees will not contribute to improve accuracy. One of the key findings observed during the experimentation process was that forest with more than 25 trees minimally contribute to the accuracy of the classifier, but can enhance the overall size of the proposed model substantially. Figure 2 illustrates a flowchart to show the overall process of the proposed method.

**Fig.2:**
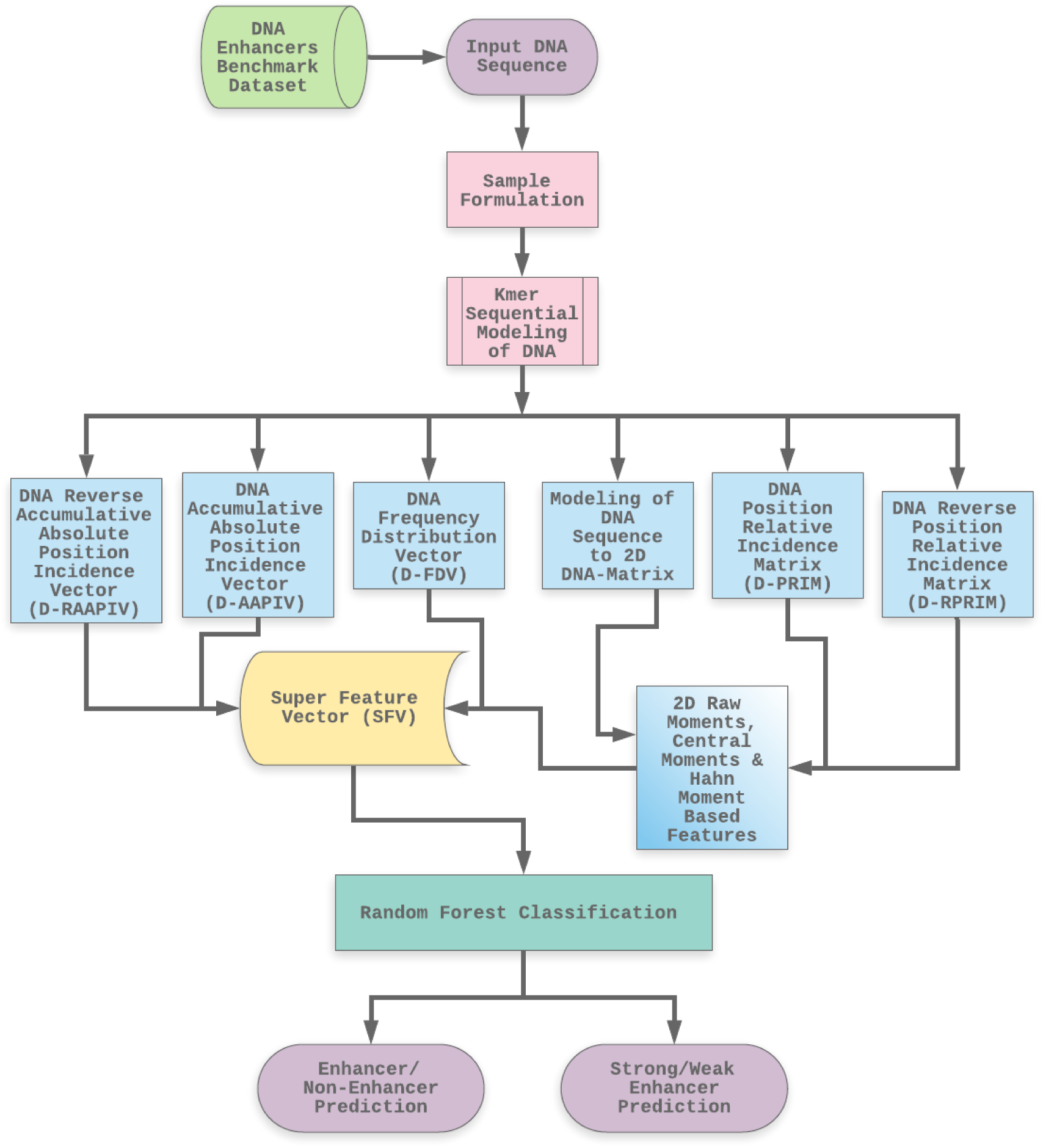
The Flowchart of the overall proposed method.

## 3. Experiments and Results

For the assessment and verifications of the model and to analyze its performance, some methods are used to evaluate them. These methods evaluate the classifiers using inspection attributes which are based on the outcomes of classification assessments and estimates.

### 3.1 Cross-validation

#### 3.1.1 k-fold Cross Validation

K-fold cross validation (KFCV) technique is most commonly used by practitioners for estimation of errors in classifications. Also known as rotation estimation, KFCV splits a dataset into ‘K’ folds which are randomly selected and are equal in size approximately. The prediction error of the fitted model is calculated by predicting the *k*^*th*^ part of the data which is dependent on other K-1 parts to fit the model. The error estimates of K from the prediction are combined together using the same procedure for each k = 1, 2, …, K.

Although the generalization performance of any classifier is mostly estimated using unbiased approximations in jackknife tests, two drawbacks exists in this test, firstly, the variance is high as estimates used in all the datasets are very similar to each other, secondly, its calculative expensive as n estimates are required to be computed, and the total number of observations to test is n in the dataset. The 5-fold and 10-fold cross validation tests are proven to be a good compromise between computational requirements and impartiality.

In the KFCV tests, the selection of ‘K’ is considered as a significant attribute. To testify errors in prediction models, cross validations (K = 5 & K = 10) tests have been used in many research studies. 5-Fold and 10-Fold tests proved to have accurate results in our proposed model and proved to be much better than state-of-the-art methods. These results are listed in Table 5 and Table 6.

### 3.2 Evaluation Parameters

The problems of binary classifications use metrics such as Accuracy (Acc), Sensitivity (Sn), Specificity (Sp) and Mathew’s Correlation Coefficient (MCC) for measuring the proposed prediction model quality and efficiency. These metrics are defined in the following eqs (24):

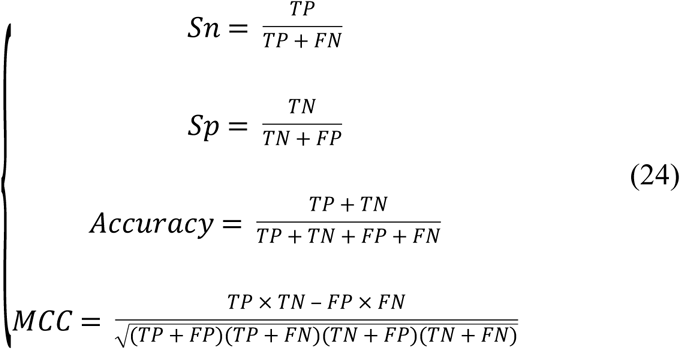

Here true-positives (TP), TN (true-negatives), FP (false-positives) and FN (false-negatives) represent the outcomes from the cross validation tests. Unfortunately, the conventional formulations from the above mentioned metrics in eqs(24) lack in intuitiveness and due to this fact, understanding these measures especially MCC, many scientists have faced difficulties. To ease this difficulty, the above conventional equations were converted by Xu [64] and Feng [65] using Chou’s four intuitive equations which used the symbols introduced by Chou [66]. The symbols that define these equations are; 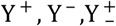 and 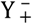. The description of these symbols is defined in Table 3.

**Table: 3.**
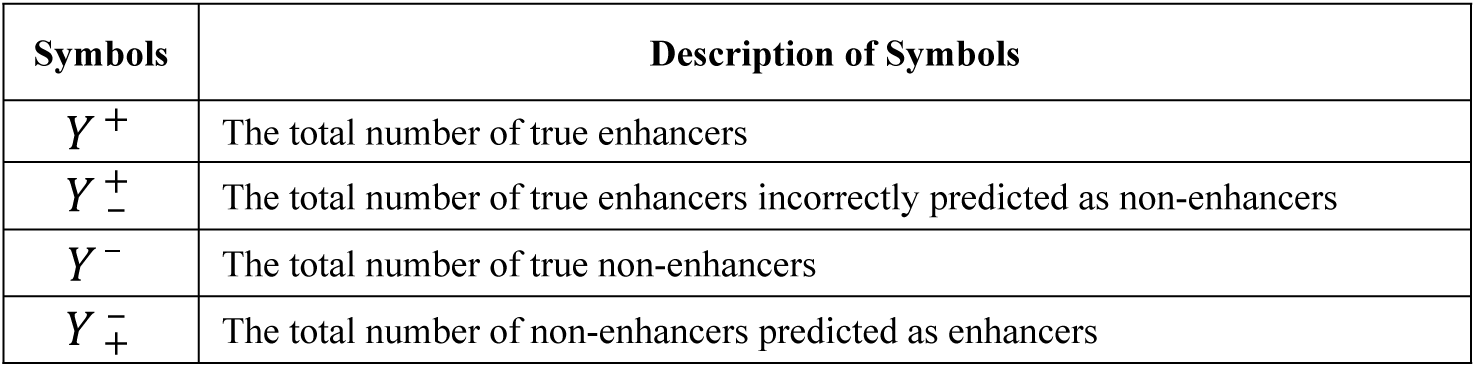
Description of symbols used to define these equations.

From the above correspondence in Table 4, we can define eqs.(25):

**Table: 4.** Description of equations used eqs. (26)

| When, | Then, | Description |
| --- | --- | --- |
| $Y_-^+ = 0$ | $Sn = 1$ | None of the enhancer is predicted as a non-enhancer |
| $Y_-^+ = Y^+$ | $Sn = 0$ | All of the enhancers were incorrectly predicted as non-enhancers |
| $Y_+^- = 0$ | $Sp = 1$ | None of the non-enhancer is incorrectly predicted as an enhancer |
| $Y_+^- = Y^-$ | $Sp = 0$ | All of the non-enhancers are incorrectly predicted as enhancers |
| $Y_-^+ + Y_+^- = 0$ | $ACC = 1,$<br>$MCC = 1$ | None of the enhancers and none of the non-enhancers were incorrectly predicted |
| $Y_-^+ = Y^+ \text{ and } Y_+^- = Y^-$ | $ACC = 0,$<br>$MCC = -1$ | All of the enhancers and all of the non-enhancers were incorrectly predicted. |
| $Y_-^+ = \frac{Y^+}{2} \text{ and } Y_+^- = \frac{Y^-}{2}$ | $ACC = 0.5,$<br>$MCC = 0$ | The overall prediction is not a better than any other random prediction outcome. |

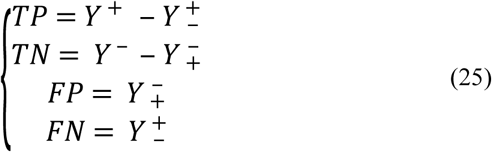

From the above correspondence in Table 4, we can define eqs.(26):

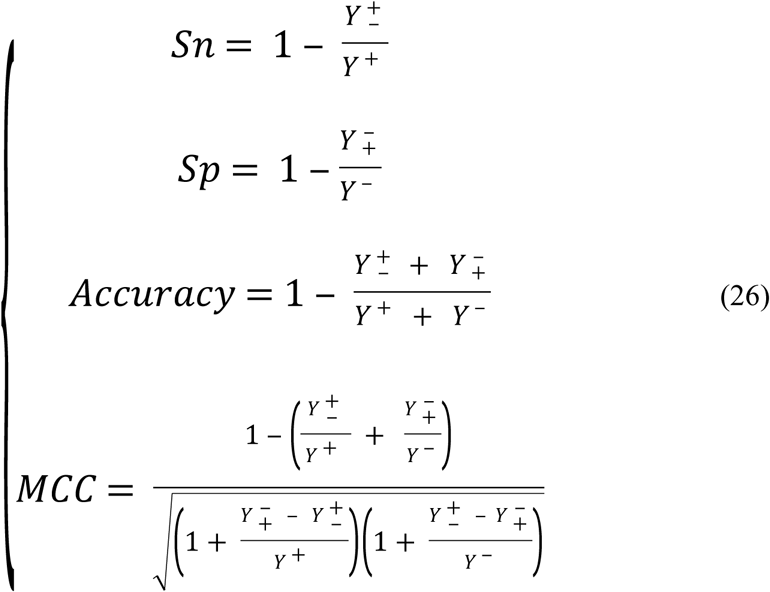

The above eqs.(26) has the same meaning as the equation (24) but it is more easy to understand and intuitive. Table 4 defines the detail description of these equations.

The set of metrics used in above Table 4 are not applicable to multi-labeled prediction models rather they are only useful for single labeled-systems. A different set of metrics exists for multi-labeled-systems which have been used by various researchers [67]–[69]. The comparison of existing classifiers with proposed method is mentioned in Table.6.

**Table: 5.** Comparison of state-of-the-art methods with the proposed method using 5-Fold Cross validation tests.

| Layer | Classifiers | Sn(%) | Sp(%) | ACC(%) | MCC |
| --- | --- | --- | --- | --- | --- |
| 1 | iEnhancer-2L | 78.09 | 75.88 | 76.89 | 0.54 |
|  | iEnhancer-2L-Hybrid | 75.33 | 80.39 | 77.86 | 0.558 |
|  | EnhancerPred | 72.57 | 73.79 | 80.82 | 0.464 |
|  | iEnhancer-EL | 75.67 | 80.39 | 78.03 | 0.561 |
|  | iEnhancer-5Step | 81.1 | 83.5 | 82.3 | 0.65 |
|  | Proposed Method | 90.77 | 92.59 | 91.68 | 0.834 |
| 2 | iEnhancer-2L | 62.21 | 61.82 | 61.93 | 0.24 |
|  | iEnhancer-2L-Hybrid | 71.02 | 60.64 | 65.83 | 0.318 |
|  | EnhancerPred | 62.67 | 61.46 | 62.06 | 0.2413 |
|  | iEnhancer-EL | 69 | 61.05 | 65.03 | 0.315 |
|  | iEnhancer-5Step | 75.3 | 60.8 | 68.1 | 0.37 |
|  | Proposed Method | 76.04 | 93.01 | 84.53 | 0.6975 |

**Table: 6.** Comparison of classifiers for predicting enhancers using 10-fold cross validations.

| Layer | Classifier | Sn(%) | Sp(%) | ACC(%) | MCC |
| --- | --- | --- | --- | --- | --- |
| 1 | KNN | 69.81 | 72.90 | 71.36 | 0.4275 |
|  | Naïve Bayes | 67.59 | 69.47 | 68.53 | 0.3712 |
|  | AdaBoost | 72.30 | 73.31 | 72.80 | 0.4569 |
|  | SVM | 70.68 | 78.43 | 74.56 | 0.4933 |
|  | PNN | 72.04 | 72.97 | 72.51 | 0.4507 |
|  | Random Forest | 86.53 | 96.90 | 91.72 | 0.8398 |
| 2 | KNN | 58.77 | 54.05 | 56.41 | 0.1285 |
|  | Naïve Bayes | 58.35 | 59.56 | 58.95 | 0.1793 |
|  | AdaBoost | 63.46 | 57.15 | 60.31 | 0.2079 |
|  | SVM | 69.94 | 55.68 | 62.80 | 0.2598 |
|  | PNN | 76.95 | 44.34 | 60.64 | 0.2261 |
|  | Random Forest | 80.49 | 93.97 | 87.23 | 0.7519 |

**Table: 7.**
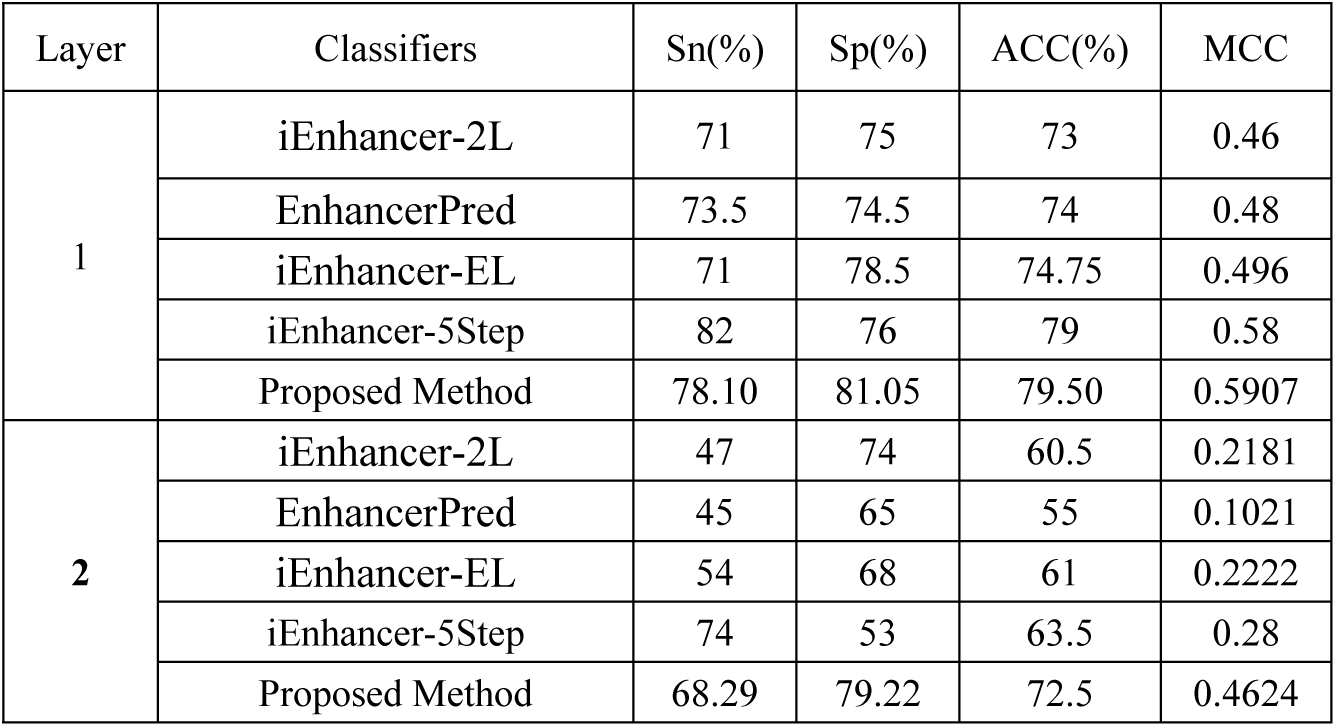
Independent tests based comparison of state-of-the-art methods with the proposed method.

| Layer | Classifiers | Sn(%) | Sp(%) | ACC(%) | MCC |
| --- | --- | --- | --- | --- | --- |
| 1 | iEnhancer-2L | 71 | 75 | 73 | 0.46 |
|  | EnhancerPred | 73.5 | 74.5 | 74 | 0.48 |
|  | iEnhancer-EL | 71 | 78.5 | 74.75 | 0.496 |
|  | iEnhancer-5Step | 82 | 76 | 79 | 0.58 |
|  | Proposed Method | 78.10 | 81.05 | 79.50 | 0.5907 |
| 2 | iEnhancer-2L | 47 | 74 | 60.5 | 0.2181 |
|  | EnhancerPred | 45 | 65 | 55 | 0.1021 |
|  | iEnhancer-EL | 54 | 68 | 61 | 0.2222 |
|  | iEnhancer-5Step | 74 | 53 | 63.5 | 0.28 |
|  | Proposed Method | 68.29 | 79.22 | 72.5 | 0.4624 |

### 3.3 Results and Discussions

The classification algorithms with their predictions results using benchmark dataset are shown in Table 5, 6 & 7. iEnhancer-EL [70] and iEnhancer-2L [26] produced better outcomes using ensemble classifiers and achieved accuracy of 78.03% and 76.89% respectively in which they were successful in predicting strong enhancers with accuracy of 65.03% and 61.93% respectively. Whereas EnhancerPred [27] achieved 80.82% accuracy and used SVMs which produced slightly better results in predicting strong enhancers with 62.06% accuracy. Similarly, iEnhancer-2L-Hybrid [71]and iEnhancer-5Step [29] improved the accuracy results with their prediction model and acquired 77.86% and 82.3% accuracies respectively with identifying the strong enhancers with 65.83% and 68.1% accuracies respectively. In contrast, 91.68% and 84.53% accuracy was achieved in predicting enhancers and their strength respectively by the currently proposed method after utilizing obscure features from statistical moments and random forest classifications using 5-Fold cross validation tests (see Table 5 and Figure.3 for ROC). Furthermore, 10-fold cross-validation test was also conducted using random forest classifier on benchmark dataset and obtained the accuracy results are listed in Table 6. The ROCs 10-fold cross-validation tests are shown in Figure.4, 5, 6, 7, 8 and Figure.9. In addition to cross validation tests, an independent test was also performed using the independent dataset. The comparison of proposed model and state-of-the-art methods using independent dataset is listed in Table Our proposed method is based on the feature sets that are evaluated using Hahn moments which are easier for the random forest based classifier to classify the feature vectors in acute time and are very efficient as compared to previous methods which were not able to produce better results on the computational cost of training and testing using classification process.

**Fig.3(a):**
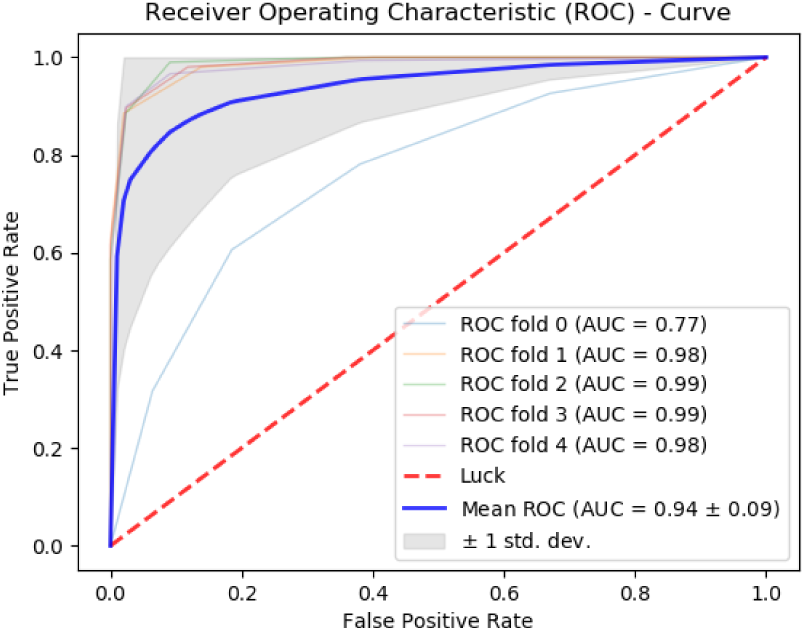
ROC Curve of 5-fold cross validation tests for enhancers.

**Fig.3(b):**
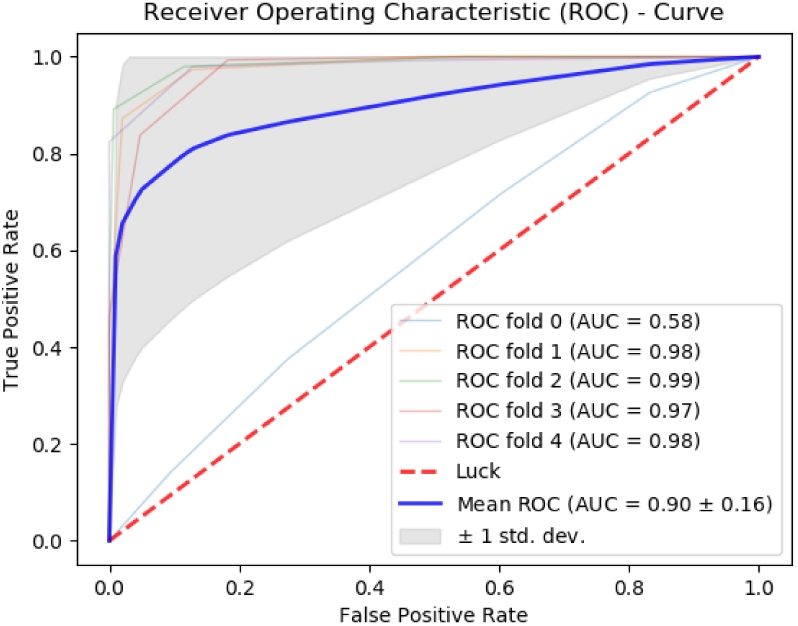
ROC Curve of 5-fold cross validation tests for enhancer strengths.

**Fig.4(a):**
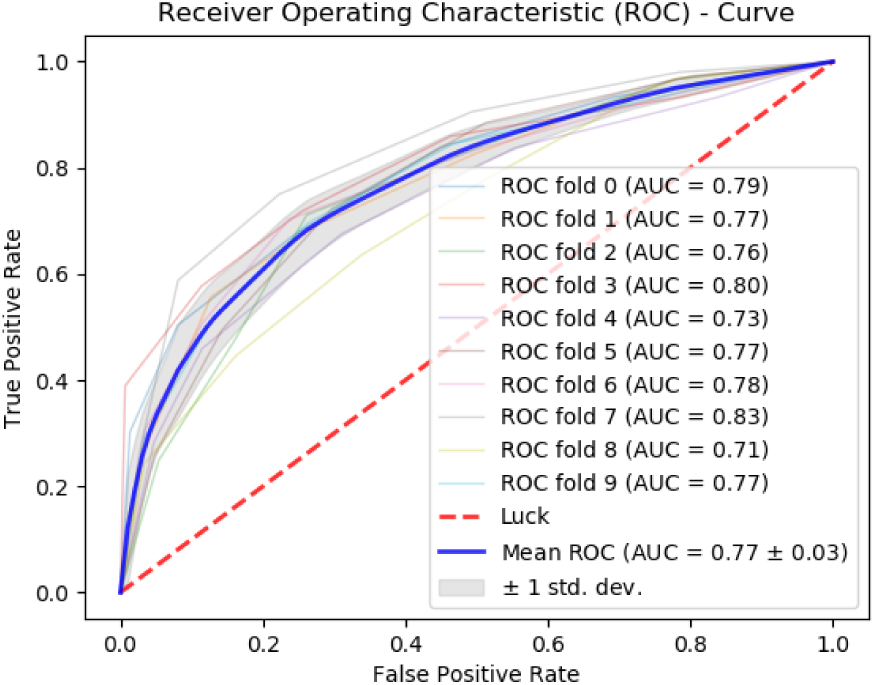
ROC Curve of 10-fold cross validation tests for enhancers using (KNN)

**Fig.4(b):**
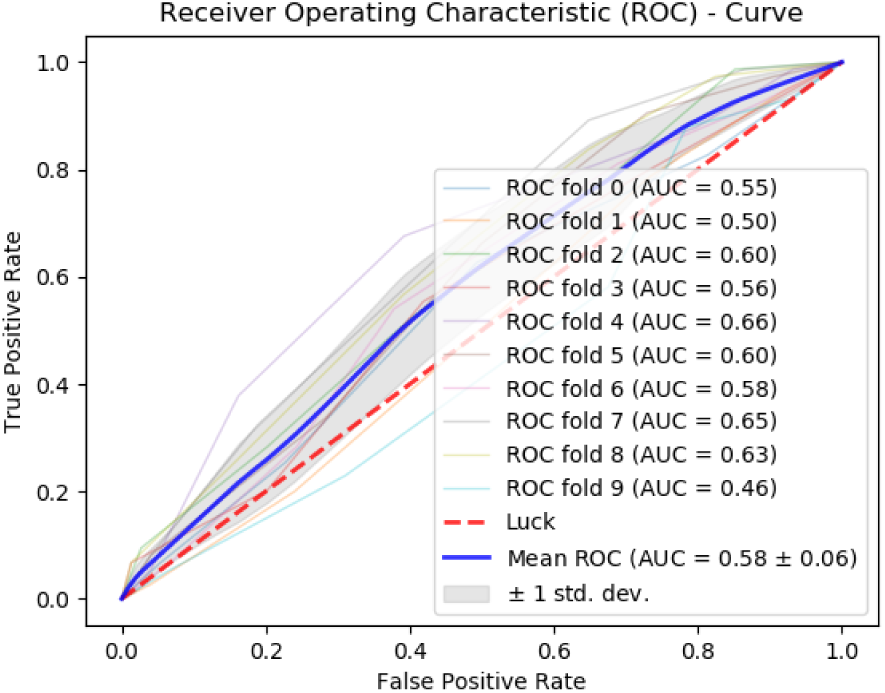
ROC Curve of 10-fold cross validation tests for enhancers strength (KNN)

**Fig.5(a):**
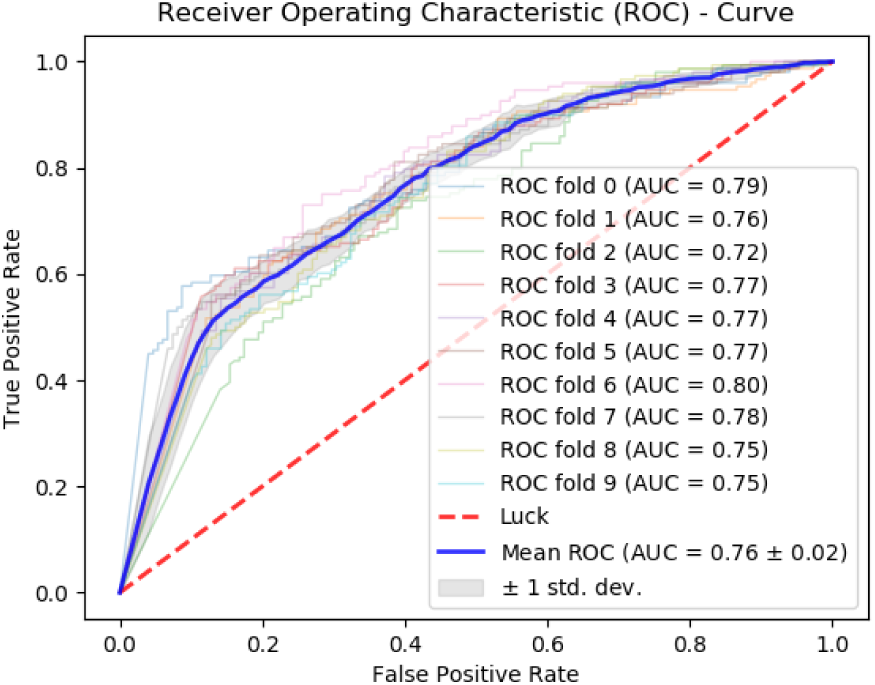
ROC Curve of 10-fold cross validation tests for enhancers using (Naïve-Bayes)

**Fig.5(b):**
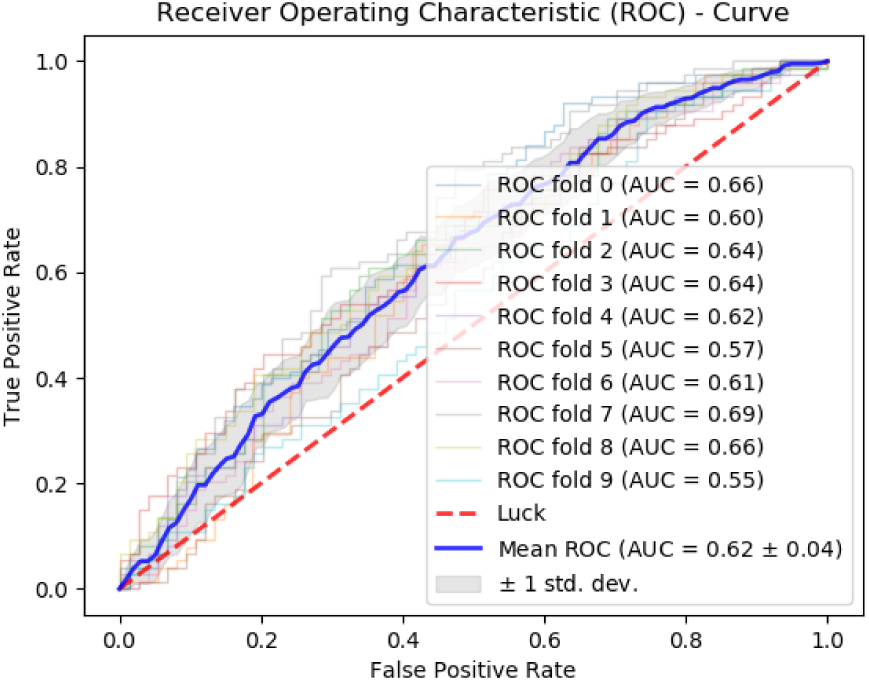
ROC Curve of 10-fold cross validation tests for enhancers strength (Naïve-Bayes)

**Fig.6(a):**
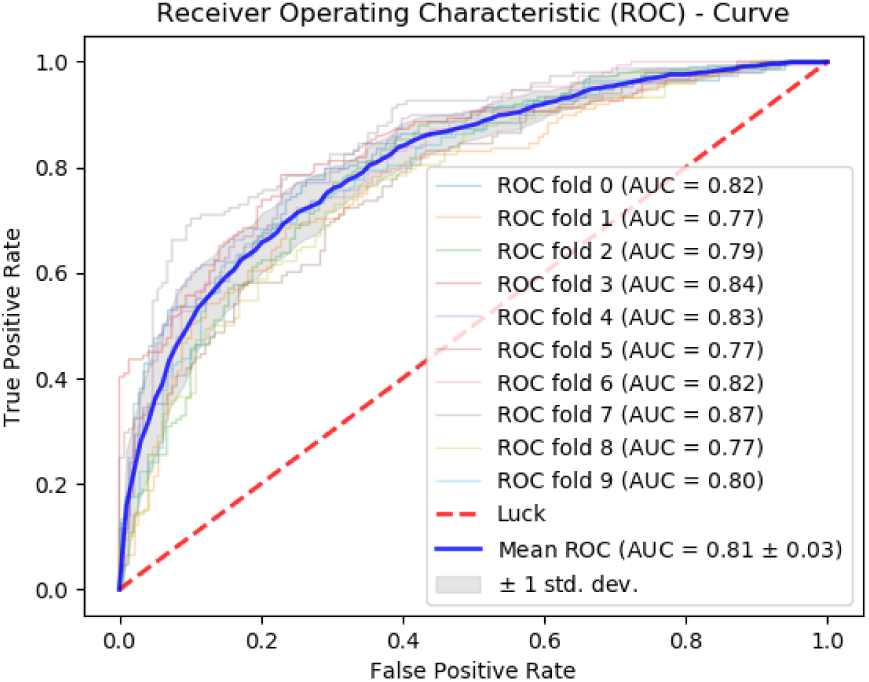
ROC Curve of 10-fold cross validation tests for enhancers using (AdaBoost)

**Fig.6(b):**
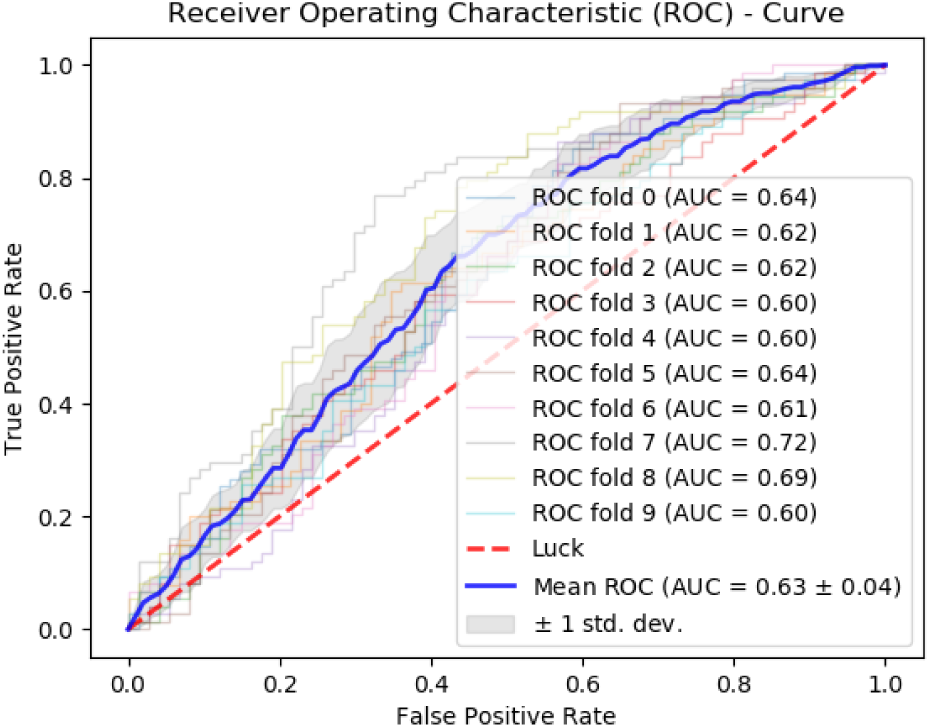
ROC Curve of 10-fold cross validation tests for enhancers strength (AdaBoost)

**Fig.7(a):**
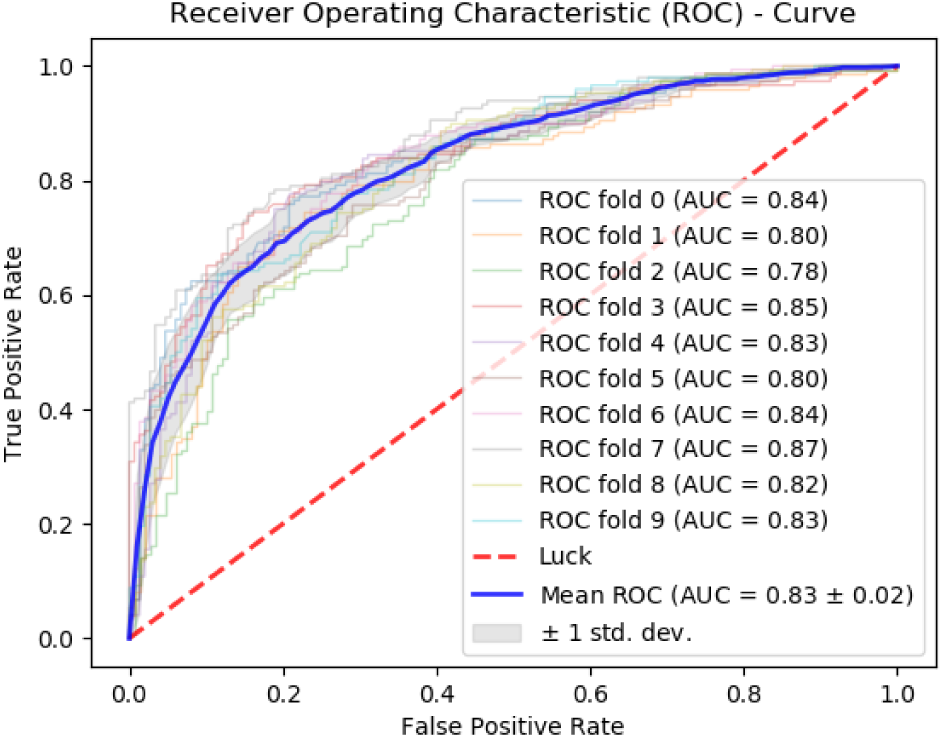
ROC Curve of 10-fold cross validation tests for enhancers using (SVM)

**Fig.7(b):**
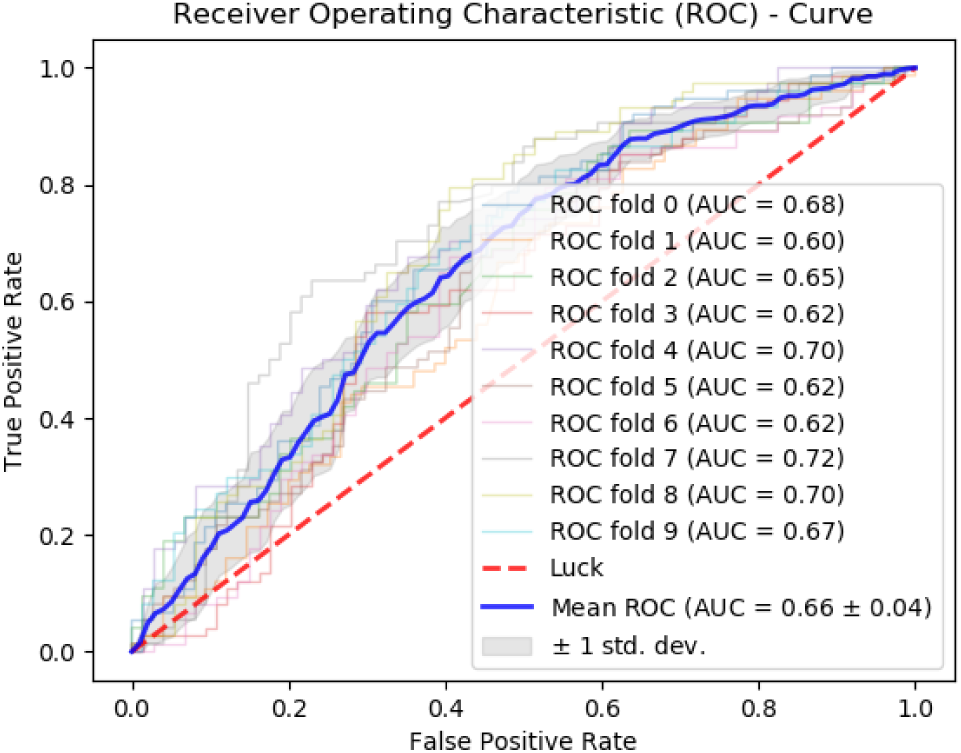
ROC Curve of 10-fold cross validation tests for enhancers strength (SVM)

**Fig.8(a):**
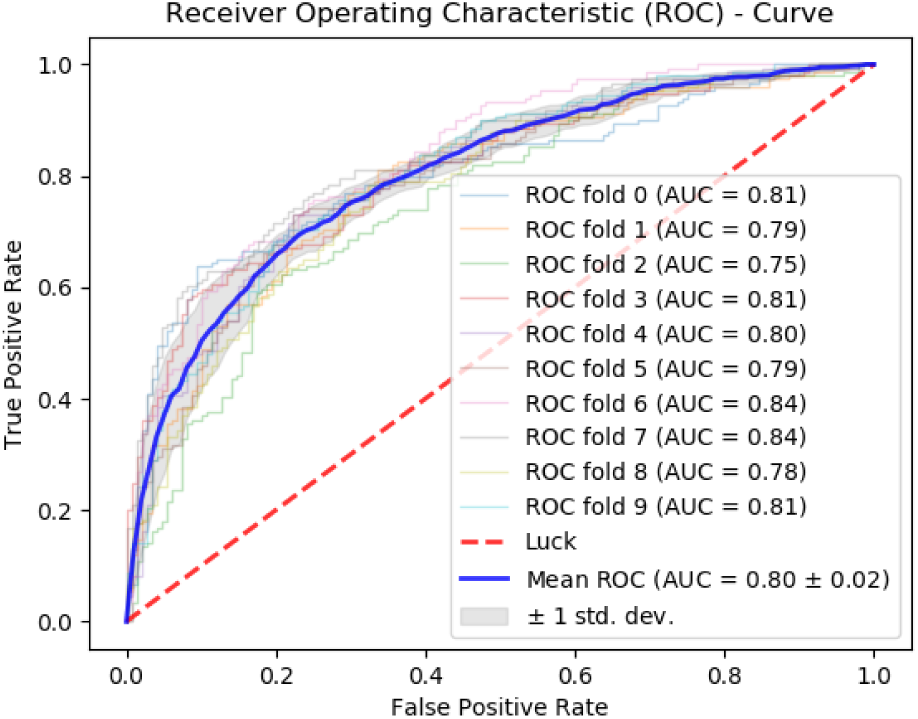
ROC Curve of 10-fold cross validation tests for enhancers using (PNN)

**Fig.8(b):**
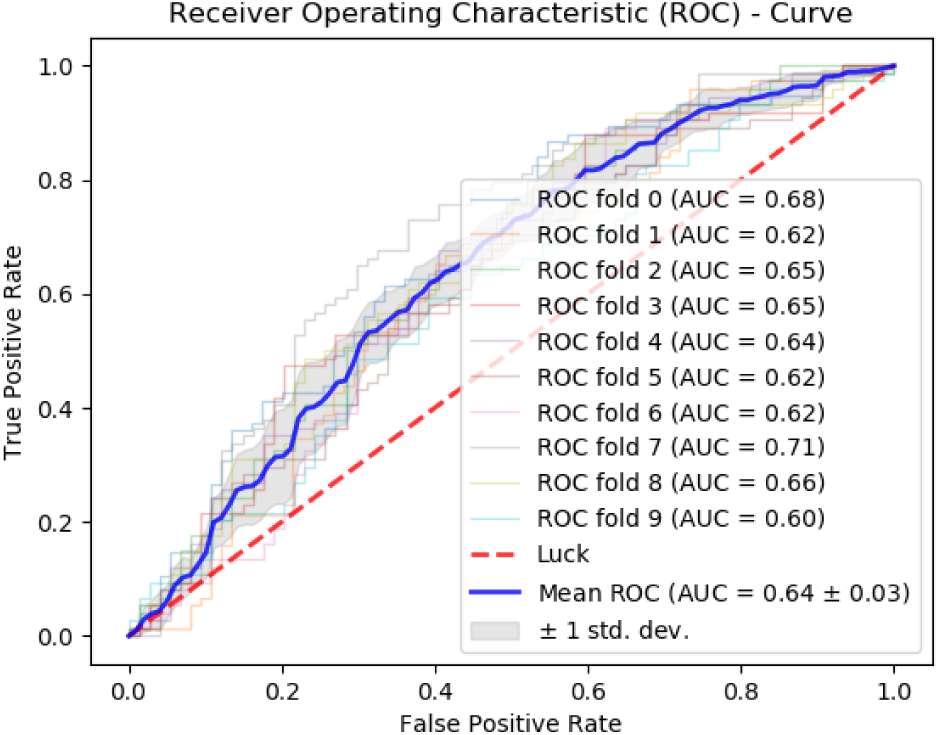
ROC Curve of 10-fold cross validation tests for enhancers strength (PNN)

**Fig.9(a):**
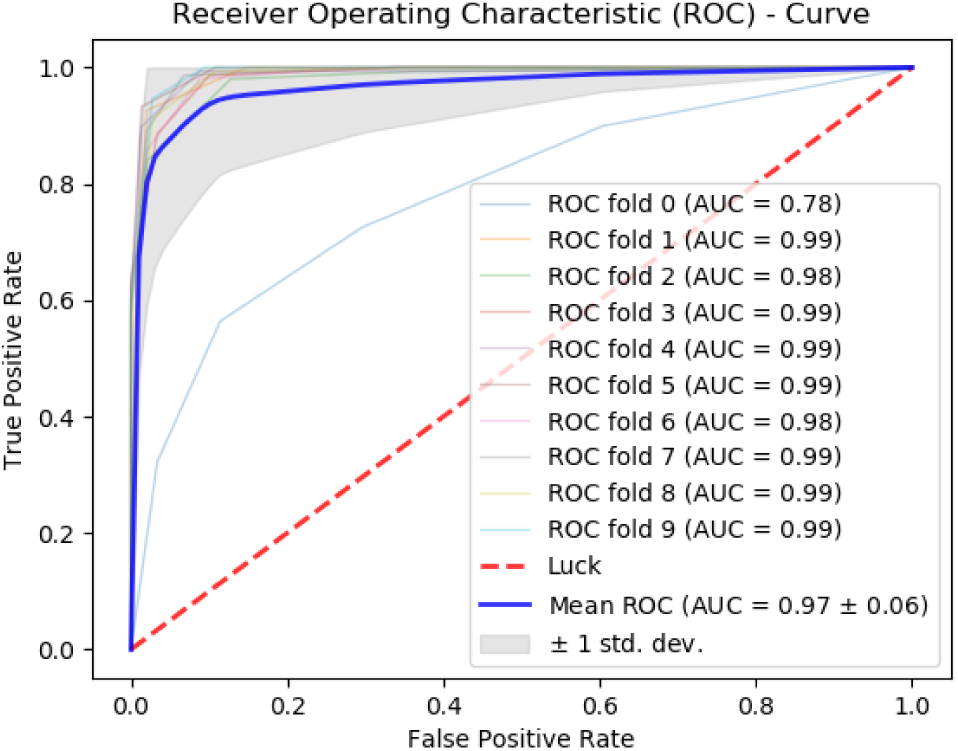
ROC Curve of 10-fold cross validation tests for enhancers using (Random Forest)

**Fig.9(b):**
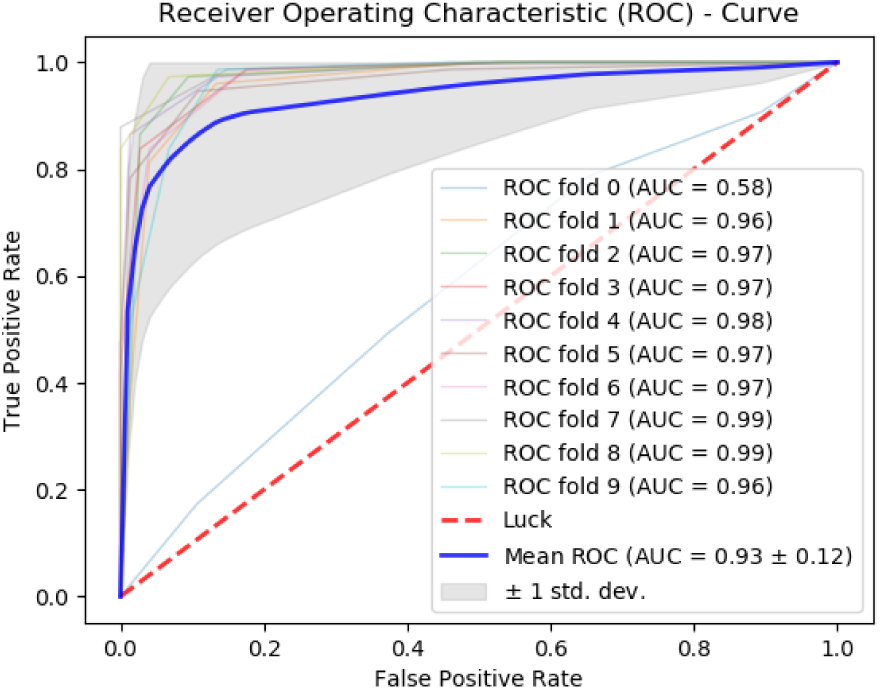
ROC Curve of 10-fold cross validation tests for enhancers strength (Random Forest)

### 3.4 Web-Server

As observed in past studies [72]–[76], the development of a web-server is highly significant and useful for building more useful prediction methodologies. Thus, efforts for a user friendly webserver have been made in past [77]–[81] to provide ease to biologists and scientists in drug discovery. The webserver which has been developed for the proposed method is freely accessible at http://www.biopred.org/enpred which is developed using Flask 1.1.1. The step-wise instructions to utilize the system are mentioned as follows:

#### 3.4.1 Step-1

Open your web browser and navigate to www.biopred.org/enpred. The first page (see Figure.10) of the webserver is **Home**, page from where you can proceed to **ReadMe, Server, Data** and **Citations** pages through provided navigation links. The **Server** tab navigates to the portal for prediction process. **Data** tab facilitates the user with links to download the benchmark datasets used during the training and testing process. Finally, the **Citations** tab redirects the user to the page where information about the relevant paper and its citation is provided. In order to perform the prediction, kindly click on the **Server** tab.

**Fig.10:**
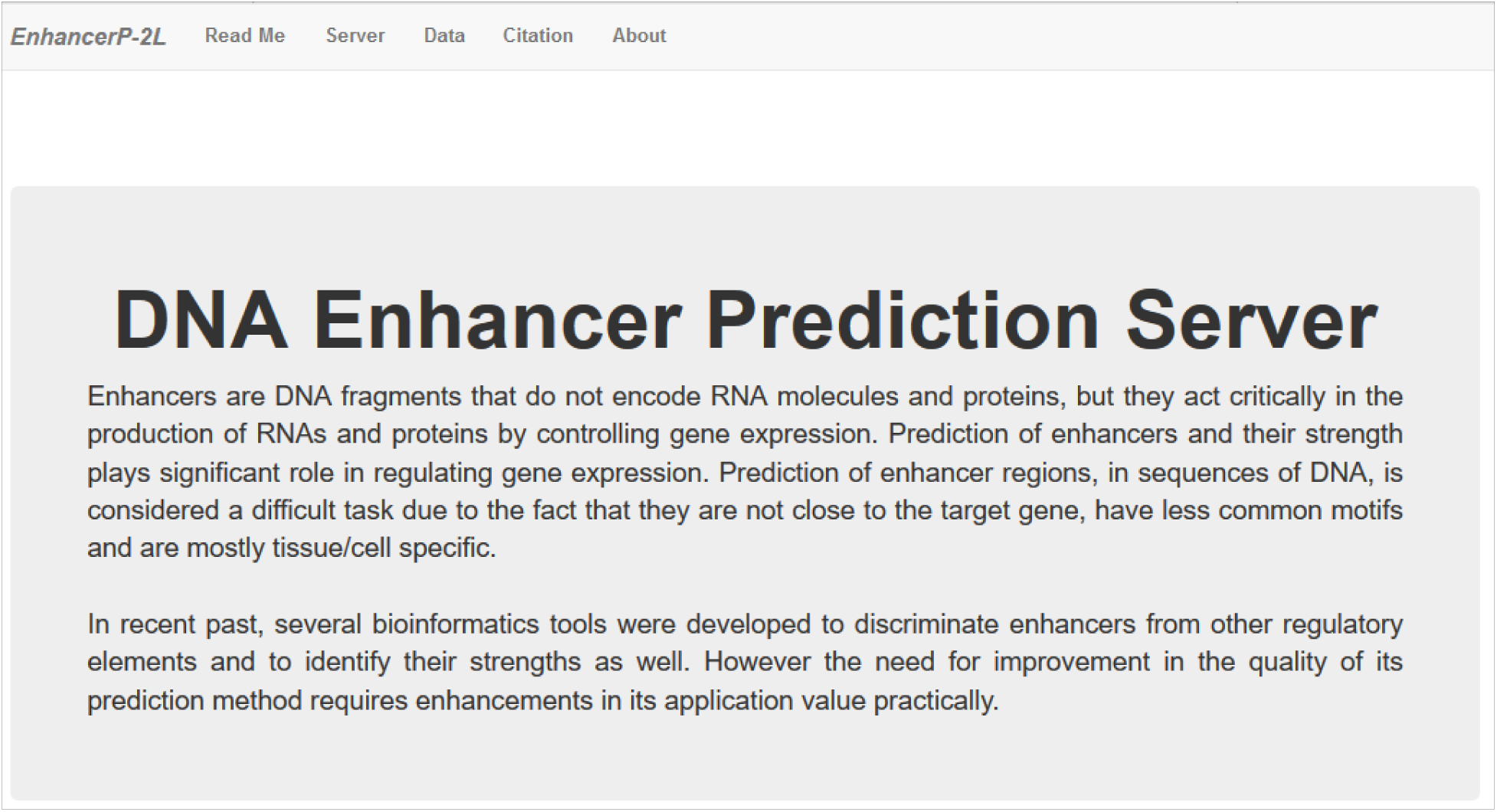
The GUI of www.biopred.org/enpred webserver for enhancer predictions.

#### 3.4.2 Step-2

On the **Server** page, the user is provided with an empty text-area where the user can input the DNA Enhancer or DNA non-Enhancer sequence for prediction. The sequence input to the webserver is required to be in FASTA format. The **Submit** button will proceed to the prediction process for the input sequence. The results of the prediction process will appear on the **Results** page. The time of the prediction process totally depends upon the length of the input sequence.

#### 3.4.3 Step-3

On the **Data** page, the user is provided with links to download the benchmark datasets for future experimentations.

#### 3.4.4 Step-4

On the **ReadMe** page, the user is provided with relevant information about the current model which includes the details about the operation algorithm used for predictions.

## 4. Conclusions

In the proposed research, an efficient model for predicting the enhancers and their strength using statistical moments and random forest classifier is developed. In recent past, many methods were proposed to predict enhancers, but our method has proved to be better in accuracy than the existing state-of-the-art methods. Our method achieved accuracies of 91.68% and 84.53% for enhancer and strong enhancer classifications on a benchmark dataset which is currently the highest and accurate classification method for prediction of enhancers and their strength.

## 5. Compliance with ethical standards

### 5.2 Ethical approval

This article does not contain any studies involved with human participants or animals performed by any of the authors.

